# Metacognitive domains are not aligned along a dimension of internal-external information source

**DOI:** 10.1101/2022.05.03.490468

**Authors:** Polina Arbuzova, Lisa K. Maurer, Elisa Filevich

**Affiliations:** Bernstein Center for Computational Neuroscience, Berlin, Germany; Berlin School of Mind and Brain, Humboldt-Universität zu Berlin, Germany; Institute of Psychology, Humboldt Universität zu Berlin, Germany; Neuromotor Behavior Laboratory, Giessen, Germany; Institute of Sport Science, Justus Liebig University, Giessen, Germany; Center for Mind, Brain and Behavior, Gießen and Marburg, Germany

**Keywords:** metacognition, domain-general, domain-specific, m-ratio

## Abstract

It is still debated whether metacognition, or the ability to monitor our own mental states, relies on mechanisms that are ‘domain-general’ (a single mechanism can account for the monitoring of any mental process) or ‘domain-specific’ (metacognition is accomplished by a collection of multiple monitoring modules, one for each cognitive domain). It has been speculated that two broad categories of metacognitive mechanisms may exist: those that monitor primarily externally-generated vs. those that monitor primarily internally-generated information.

To test this proposed division, we measured metacognitive performance (using m-ratio, a signal detection theoretical measure) in four tasks that could be ranked along an internal-external axis of the source of information, namely memory, motor, visuomotor and visual tasks. We found correlations between m-ratios in visuomotor and motor tasks only; but no correlations between m-ratios in visual and visuomotor tasks, or between motor and memory tasks.

While we found no correlation in metacognitive ability between visual and memory tasks, and a positive correlation between visuomotor and motor tasks, we found no evidence for a correlation between motor and memory tasks. This pattern of correlations does not support the grouping of domains based on whether the source of information is primarily internal or external. We suggest that other groupings could be more reflective of the nature of metacognition and discuss the need to consider other non-domain task-features when using correlations as a way to test the underlying shared mechanisms between domains.

## Introduction

The human brain processes a vast variety of information, and is capable of perceiving both the outside of the world (through vision, audition, or olfaction), and bodily signals (through interoception and proprioception). Additionally, it can process information generated internally, when retrieving information stored in memory or when experiencing emotions. Each of these kinds of information can be said to constitute a cognitive domain (Spunt & Adolphs, 2017). The human brain is also capable of introspection, or metacognition: It can form second-order representations of its own cognitive processes (Fleming & Dolan, 2012; Rouault et al., 2018). Intuitively, these second-order metacognitive representations might be less differentiated into domains as compared to first-order cognitive processing (Fleming, 2020), because the former are higher in the cognitive hierarchy than the latter (Brown et al., 2019). Despite its intuitive appeal, this is an empirical question (Rouault et al., 2018): To what extent are metacognitive mechanisms the same across several cognitive domains? Further, which factors might allow the same metacognitive mechanisms to monitor different cognitive domains?

Experimentally, metacognition is typically operationalised by asking participants to rate their confidence in their own perceptual or cognitive decisions, on each one of often hundreds of trials (Fleming and Lau 2014). Measures of metacognitive ability in these paradigms quantify how well high confidence responses track correct decisions. Then, one common way to test whether a single mechanism underlies metacognitive monitoring across different domains involves asking participants to monitor their performance in different domains and measure correlations in metacognitive ability across domains: If measures of metacognitive performance in two domains are not correlated, this indicates that they have different underlying mechanisms (and the presence of a correlation is normally interpreted as a sign that underlying mechanisms at least partially overlap). Several studies have followed this logic to measure correlations between metacognitive ability across different domains. Overall, results are mixed: Some studies revealed positive correlations between metacognitive ability across different perceptual domains (Faivre et al., 2018; Samaha & Postle, 2017; Song et al., 2011) and across vision and memory domains (Mazancieux et al., 2018; McCurdy et al., 2013); whereas other studies revealed dissociations between domains (memory vs vision: Baird et al., 2013, 2015; Fitzgerald et al., 2017; Morales et al., 2018, nociception vs vision and nociception vs thermoception: Beck et al., 2019; tactile perception vs interoception: Garfinkel et al., 2016; auditory perception vs interoception: Legrand et al., 2022). In a meta-analysis, Rouault et al (2018) considered two categories: Studies that tested two or more perceptual tasks, and those that tested one memory and one perceptual task. This revealed a positive meta-analytic correlation for studies including perceptual tasks only, but none for studies including memory and perceptual tasks. Further, a meta-analysis of neural correlates of metacognitive judgements (Vaccaro and Fleming, 2018) suggested a partially distinct network for perceptual and for memory domains: While insula, the lateral and posterior medial prefrontal cortex (PFC) are involved in both, left dorsolateral PFC and bilateral parts of the parahippocampal cortex are activated specifically in metamemory tasks.

To map the relationships between domains at the metacognitive level, one (inefficient) approach would be to test metacognitive ability in all possible domains and examine pairwise correlations between them. A more efficient way to understand these relationships would be to identify general principles that describe them. The results of the two meta-analyses mentioned above (Rouault et al., 2018; Vaccaro & Fleming, 2018) suggest one potential general principle: Two broad, separate metacognitive domains might exist to monitor primarily externally-generated (for example, sensory information) vs. primarily internally-generated information (for example, memory and motor commands) (Fleming et al., 2014; Rouault et al., 2018). Because most studies probed domain-generality by asking participants to complete perceptual (mostly visual) and memory tasks, it is impossible to determine whether the general distinction between internally- and externally-generated is valid, or whether it applies only to those perceptual and memory processes that were tested. To explicitly test this proposed general principle of organization of metacognitive processes, it is necessary to test metacognition in domains beyond vision and memory. Here, we capitalized on the unique features of the monitoring of voluntary movement: Unlike perception and memory, which are clearly primarily externally- and internally-generated, voluntary movements are associated with both kinds of sources of information that participants might concurrently monitor to make metacognitive judgments. Both internal efferent motor commands, as well as external afferent signals (including vision, audition and proprioception) (Miall & Wolpert, 1996) may be informative for metacognitive representations. We used a visuomotor metacognitive task, where externally-generated information (visual and proprioceptive) and internally-generated information (motor commands) are available for monitoring; as well as a motor task, where visual information is not available and monitoring is therefore less reliant on externally-generated information. In that way, small modifications of a single metacognitive task allowed us to control the relative availability of internally- and externally-generated information. We examined correlations in metacognitive ability between these two variations of the task, and between them and a visual and a memory task, which represent the extremes of the hypothesized external-internal principle of organization.

Our working hypothesis was that, if the internal-external axis is useful in guiding the distinction between domains of metacognitive monitoring, the pattern of correlations of individual metacognitive ability would correspond to the distance between each pair of tasks, on the basis of their expected positions along the axis. We made five predictions. First, we expected the lowest correlation (or at the extreme, none at all) between the visual and memory tasks. Second, we expected to find a positive correlation between the (visuo)motor and memory tasks, because they require participants to monitor internally-generated information. Third, we expected a positive correlation between visuomotor and visual tasks because of the shared visual (external) information. Fourth, we expected a positive correlation between visual and motor tasks because they both share external (visual and proprioceptive) sources of information. And finally, we expected a positive correlation between visuomotor and motor tasks because of both the shared external (visual and proprioceptive) and internal (motor command) sources of information. The first three hypotheses were pre-registered, and the final two are based on our previous study (Arbuzova et al., 2021) and follow the same logic as the others.

## Methods

The study and analyses were pre-registered (https://osf.io/6u3sj/). The data, experimental and analysis scripts are available online (https://osf.io/bwkfp/). Unless stated otherwise, we followed our pre-registration plan.

### Participants

Forty participants completed the study (21 female and 19 male, mean age 26.28 years, SD = 3.76). The sample size was based on our previous study (Arbuzova et al., 2021). All participants were right-handed, had normal or corrected-to-normal vision, and no history of neurological or psychiatric disorders (as per self-report). Participants were recruited using the university’s online recruitment platform and social media, and were reimbursed for their time and effort at the rate of 8€/h. Participants were naive regarding the hypotheses of the study. The experiment was approved by the ethics committee of the Humboldt-Universität zu Berlin.

### Apparatus

All stimuli were displayed on an LCD monitor (2560 × 1440 pixels, 61 cm × 34.5 cm, refresh rate 60 Hz), placed approximately 50-60 cm away from the participant. Participants used a response box (Black Box ToolKit, York, UK) to make a discrimination decision and rate confidence. For the visuomotor task, participants used a custom-made manipulandum to control and ‘throw’ a virtual ball on the monitor in order to hit a virtual target standing behind a virtual obstacle. The manipulandum was placed under the participant’s forearm, and consisted of a horizontal metal bar that pivoted around a vertical axis under the elbow. A goniometer (Novotechnik RFC4800 Model 600, 12 bit resolution, corresponding to at least 0.1° precision) at this joint was used to measure the angle of the metal bar. An electrical switch at the tip of the metal bar under participants fingers (similar to a touch sensor) was used to control the release of the ball. A Labjack T7 (LabJack Corporation, Lakewood, CO) data acquisition device transferred analogue data from the goniometer and the electric switch with a sampling rate of 1000 Hz.

### Stimuli and procedure

Each participant completed four tasks (at least 180 trials each; in the Memory task, this number varied and could be up to 196 trials). Each trial in all four tasks followed the same basic structure (Figure 1): A two-alternative forced-choice (2AFC) discrimination judgement followed by a confidence rating on a scale from 1-4, labelled as “very unsure”, “unsure”, “sure”, and “very sure” in German. The 2AFC and confidence decisions were self-paced in the visual and memory task; whereas in the visuomotor and motor tasks, the confidence rating response or next trial started after 10 s (due to limited capacity of the movement recording buffer), but we excluded trials with response times longer than 8 seconds, described below.

**Figure 1.**
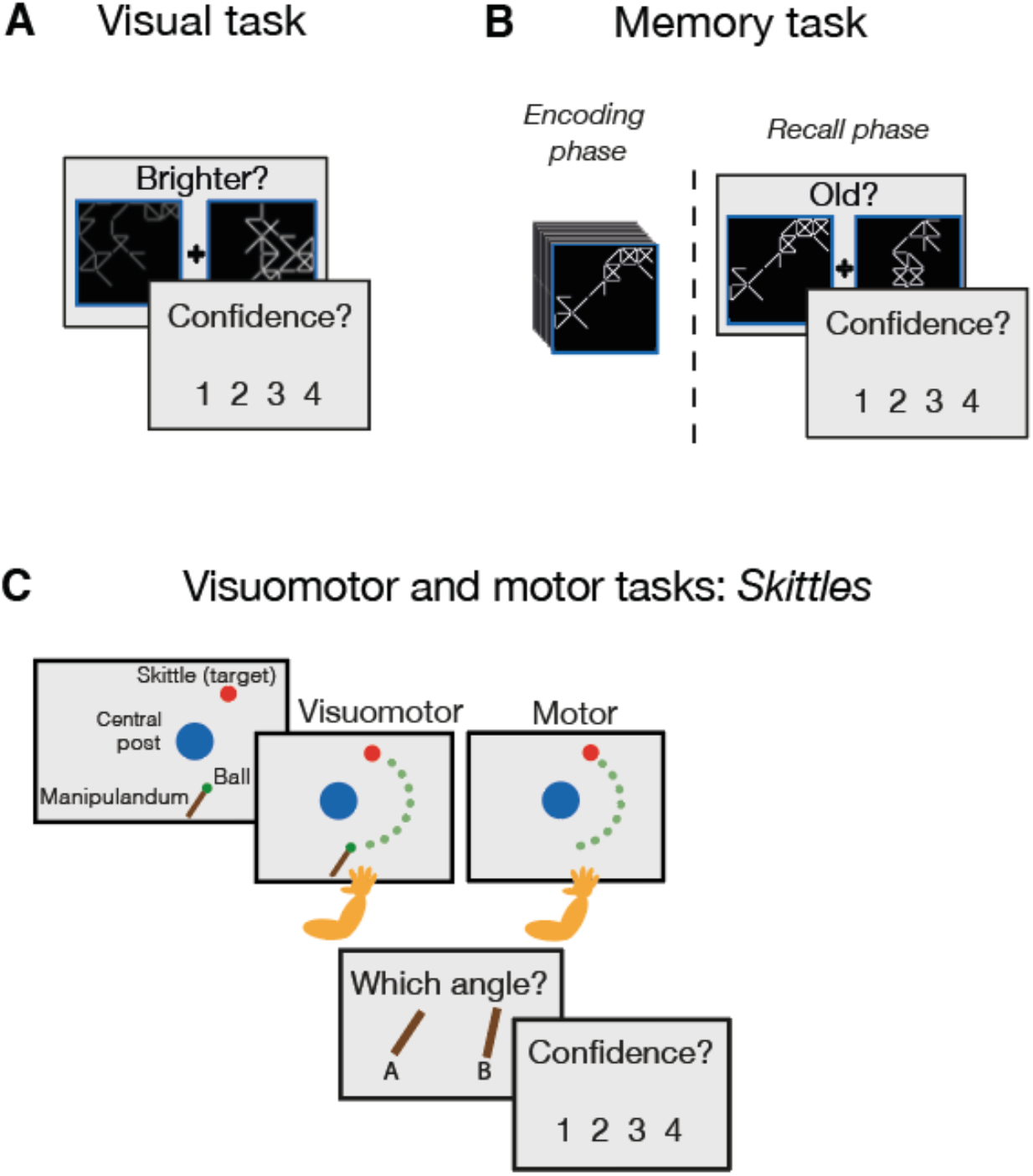
Experimental paradigms. For all tasks, each trial consisted of a 2AFC distrimination decision, followed by a confidence rating on a four-point scale **A. Visual task**. Participants briefly saw two abstract figures consisting of lines on a grid, and responded with a keypress which side they thought had the brighter lines. **B. Memory task**. Participants first memorized a number of abstract lines stimuli in the encoding phase. Then, in the recall phase, they saw two stimuli side by side, one of which was included in the encoding set and the other one was not, and responded with a keypress which side contained a figure that they had seen during the previous encoding phase. **C. Visuomotor and motor tasks (*Skittles* task)**. In this semi-virtual throwing task, participants made an accelerating movement to throw a virtual ball around a post. They saw the scene from a bird’s-eye view. In the visuomotor task, they saw a bar that corresponded to the manipulandum and their arm on the screen, whereas in the motor task, they did not. After each throw, participants responded with a keypress which of two bars best represented the angle of their arm at the moment of ball release.

The difficulty in all tasks was controlled with an online staircase (see further details below). The visual and memory tasks were adapted from Morales et al. (2018). The visuomotor and motor tasks were based on the metacognitive version of the Skittles task (Müller & Sternad, 2004) and adapted from our previous work (“angles task” in Arbuzova et al., 2021). The order of the tasks (visual, memory and the two versions of the Skittles tasks) was counterbalanced between participants. The order of visuomotor and motor trials in the Skittles task was pseudorandomized. In total, each experimental session lasted approximately 2 hours.

### Visual Task (Figure 1.A)

Each trial of the visual task started with a fixation cross. Two stimuli (sets of straight white lines, pseudorandomly placed on a black background and connecting vertices of an invisible 6×6 grid) appeared within blue square placeholders at either side of the fixation cross for 100 ms. Participants used the left/right keys on the keypad to indicate which of the two sides contained the brighter set of lines. The placeholders became thicker for 500 ms to provide feedback for the chosen side. No feedback about the accuracy of the choice was given during the main part of the task. The intertrial interval was 500 ms. The difficulty in the visual task was controlled with a 2-down-1-up online staircase on the line luminance.

To allow participants to familiarize themselves with the task, they first performed 12 trials with feedback about their accuracy (but no confidence ratings) and then 12 trials (or less, if participants indicated that they understood the task) with confidence ratings, where feedback about the accuracy of the first-order response appeared after the confidence response. Finally participants completed 40 trials with no feedback and no confidence ratings, before starting the main task. The difficulty of the first-order task in this last set of 40 trials was controlled with a staircase, to find an adequate starting point for the difficulty in the main task.

### Memory Task (Figure 1.B)

In the memory task, participants observed a number of stimuli, presented one after the other, in an encoding phase. As in the visual task, each stimulus consisted of connected white lines placed pseudorandomly. The number of stimuli presented on each encoding phase ranged between 2 and 17 (with a mean of 10 across all participants and SD of 1.06) and it was determined by an online staircase. Then, on each trial of a recall phase, two stimuli were presented at either side of the fixation cross for 2000 ms: The target stimulus that had been presented during the encoding phase, and a lure that had not been presented before. None of the lure stimuli had ever been targets in different encoding phases or vice versa.

The difficulty of the first-order memory task was staircased by changing the number of the to-be-memorized items presented in the encoding phase, increasing or decreasing by 1 if participants responded correctly to at least 80%, or less than 60% of the trials during the recall phase, respectively, averaged over the current and all previous blocks. This resulted in a different number of trials per block for different participants, and consequently, in a different total number of trials for each participant.

Before the main part of the task, participants completed a series of training trials. The training included 15 trials (three encoding-recall blocks) with only the first-order task and feedback about the accuracy of the discrimination response, followed by 18 trials (two encoding-recall blocks) with a confidence rating after each discrimination response and feedback about its accuracy. Finally, participants completed five encoding-recall blocks with no feedback about the first-order task and no confidence ratings. The number of stimuli in each block was adjusted using the staircasing method as described above, with nine stimuli in the first block. The final number of items was used as a starting number of the set size of the first encoding phase of the main part of the task.

### Visuomotor Task (Figure 1.C)

The visuomotor Skittles task was a semi-virtual ball throwing task. On each trial participants swung their forearm with the manipulandum on the horizontal plane and lifted their index finger to release a virtual ball. This setup allowed us to use a naturalistic movement, while also restricting its degrees of freedom. The ball trajectory was fully determined by two parameters: the angular velocity and the angle at the point of ball release (for details about the full model, see Müller & Sternad, 2004, and Sternad et al., 2011). During the ball throw, participants saw the scene from above on the monitor, consisting of a bar that represented their moving arm on the manipulandum, the ball at the distal end of the bar, the target, and the obstacle. The flying ball appeared on the screen for 1 s after the point of release. After each throw, participants chose which of two tilted bars displayed on the screen best represented the angle of their arm at the point of ball release. The difficulty of the Skittles task was controlled with a 2-down-1-up online staircase on the angle difference between the two alternative bar positions presented.

Before the main part of each task, participants had a chance to gradually familiarize themselves with the Skittles task. They first performed 8 ball throws, to get used to the mechanics of the virtual ball game. Then the 2AFC task was introduced and participants did 16 trials (8 for each visuomotor and 8 for motor condition), with trial-wise feedback about the accuracy of their response. After that, participants also completed 8 trials (4 of each condition) with the 2AFC task and the confidence rating (also with feedback). Finally, to find the optimal starting point for the main experiment, participants completed 96 trials (48 of each condition) with a 2-down-1-up online staircase on the line luminance, with the 2AFC task and without the confidence ratings. This part did not contain any feedback about task performance.

### Motor Task

The motor Skittles task was exactly like its visuomotor counterpart, but differed in that the critical visual information (namely, the bar representing their arm) was not visible either before, during or after the ball throw.

## Data analysis

### Metacognitive efficiency

To quantify metacognitive performance in the different tasks, we used the type II SDT-based measure *m-ratio* (*meta-d’/d’*) (Maniscalco & Lau, 2012). *M-ratio* reflects type II performance *(meta-d’*) normalized by type I performance (*d’*) and is also referred to as metacognitive efficiency. Further in this article, we also reserve this term for *m-ratios*. To estimate *m-ratios*, we used the R package metaSDT (Craddock, 2018) with the maximum likelihood fitting procedure and Broyden– Fletcher–Goldfarb–Shanno optimization algorithm.

### Robust correlations

We estimated correlations with the Robust Correlations toolbox (Pernet et al., 2013), running on Matlab (Mathworks, Natick, MA). Specifically, we used the skipped correlation method, which removes bivariate outliers as per the box-plot rule. We followed the recommendations (Pernet et al., 2013) and used Pearson’s or Spearman’s r, depending on the data distribution. To visualize correlations, we used the *lmodel2* package in R (Legendre & Oksanen, 2018) to calculate the line of the best fit with the major axis regression method, which minimizes the perpendicular distance from a point to the line.

### Bayesian statistics

We computed Bayes factors (BF_10_) to quantify the evidence for or against each correlation coefficient being different from 0. We used the BayesFactor package in R (Morey & Rouder, 2018) to estimate BF_10_ values for Pearson’s r, and neatStats package in R for non-parametric Spearman’s r (Lukács, 2021; van Doorn et al., 2020).

### Exclusion criteria

We pre-registered, and followed, the same exclusion criteria as in Arbuzova et al. (2021). We excluded trials with response times shorter than 0.2 s and longer than 8 s (<1%). We excluded results from individual tasks if the accuracy was below 60% or above 80%. To ensure that we obtained stable m-ratio estimates, we excluded data from individual participants and tasks if the type I or type II false alarms or hit rates, after splitting confidence ratings into “low” (1 or 2) and “high” (3 or 4) for a participant in a given task was smaller than 0.05 or larger than 0.95 (Bor et al., 2017). Since this affected the number of participants included in each correlation pair, we state the effective sample size for each correlation separately.

## Results

### Mean differences between tasks

First, we compared metacognitive efficiency, *m-ratio* (Figure 2.C) between tasks. A one way repeated-measures ANOVA revealed a main effect of task on the *m-ratios* (ANOVA: F(3, 54) = 12.13, p < .001, mean (M) *m-ratios* and standard deviations (SD) per condition: visuomotor: M = 0.65 (0.53), motor: M = 0.66 (0.51), visual: M = 0.84 (0.46), memory: M = 1.38 (0.34)). Bonferroni-corrected, post-hoc pairwise t-tests confirmed that *m-ratios* in the memory task higher than in all other three tasks (all corrected p-values ≤ 0.004). This result is in line earlier findings from Morales et al (2018) and might reflect post-decisional processes after the type I decision (Moreira et al., 2018; Pleskac & Busemeyer, 2010). Speculatively, memory is more susceptible for them than other domains because reflecting upon one’s recollections is an activity that we often engage in real life (for example, when having to communicate one’s recollection) and might be more ecologically valid than in other domains.

### Correlations between tasks

For our main analyses, we ran pairwise correlations between *m-ratios* obtained for each task, resulting in six correlations.

#### Memory vs (visuo)motor tasks

First, we examined relationships between the modalities on the left side of the proposed internal-external spectrum for information monitoring, namely the memory and (visuo)motor tasks. All three tasks involve the monitoring of internally-generated information. The correlation between m-ratios in the memory and in the motor task was significant, but contrary to the predictions of the internal-external axis hypothesis, it was negative (Spearman’s r = -0.56, confidence interval (CI) [-0.81 -0.51], n = 24, BF_10_ = 6.47). The correlation between *m-ratios* in visuomotor and memory modalities was not significant, Pearson’s r = -0.44, CI: [-0.77 -0.02], n = 18, BF_10_ = 1.35, although note that the CI was skewed heavily towards negative values, suggesting that there might be a negative correlation that our study did not have enough power to detect.

#### Visual vs. memory tasks

We then examined the relationships between metacognitive ability between the two extremes of the proposed internal-external axis, namely the visual and memory tasks. In line with previous literature (Baird et al., 2013, 2014, 2015; Fitzgerald et al., 2017) and with the previous study from which we derived our memory tasks (Morales et al., 2018), we found no correlation between metacognitive efficiency between the visual and in memory tasks (Pearson’s r = -0.17, CI: [-0.53 0.17], n = 24, BF_10_ = 0.35) (but see Mazancieux et al., 2018; McCurdy et al., 2013; Samaha &Postle, 2017; Song et al., 2011 for conflicting results).

#### (Visuo)motor vs. visual tasks

Finally, we examined correlation pairs in *m-ratios* between visuomotor, motor and visual conditions. We found a positive correlation between *m-ratios* in the motor and the visuomotor conditions, (Spearman’s r = 0.57, CI: [0.24 077], n = 30, BF_10_ = 18.29), as well as a positive and significant correlation between *m-ratios* in the motor and visual tasks (Pearson’s r = 0.46, CI: [0.11 0.76], n = 30, BF_10_ = 4.45). Both these results are in line with our previous work (Arbuzova et al., 2021). We found no significant correlation between the visual and the visuomotor tasks (Spearman’s r = 0.27, CI [-0.11 0.65], n = 28, BF_10_ = 0.56), unlike our previous findings from Arbuzova et al. (2021).

Results for levels of *d’, m-ratios* and mean confidence ratings across tasks are provided in Supplementary Materials.

## Discussion

In this study, we tested the hypothesised distinction between metacognitive mechanisms for monitoring of internally and externally generated sources of information (Fleming et al., 2014; Rouault et al., 2018). We considered four different metacognitive tasks (memory, motor, visuomotor, visual) that fall on this hypothesised internal-external axis. We take metacognitive monitoring in the visual task to be strongly reliant on externally-generated information; and in the memory task to be strongly reliant on internally-generated information. Also, we consider the two variations of the motor metacognitive task to lie between these extremes.

If this internal-external categorization is a valid one, it should apply to all types of metacognitive monitoring, and not just to the two specific cases of visual and memory monitoring from which it was deduced. Therefore, we expected metacognitive efficiency, measured through *m-ratios*, to be more strongly associated between neighbouring modalities and less so between modalities lying in the extremes, resulting in a given pattern of pairwise correlations (Figure 2A). However, our data do not support that — most strikingly, we found a negative correlation between metacognitive efficiency in memory and motor domains. Thus, the pattern of correlations we obtained did not match the pattern that would be expected if the internal-external axis of metacognitive monitoring were true.

**Figure 2.**
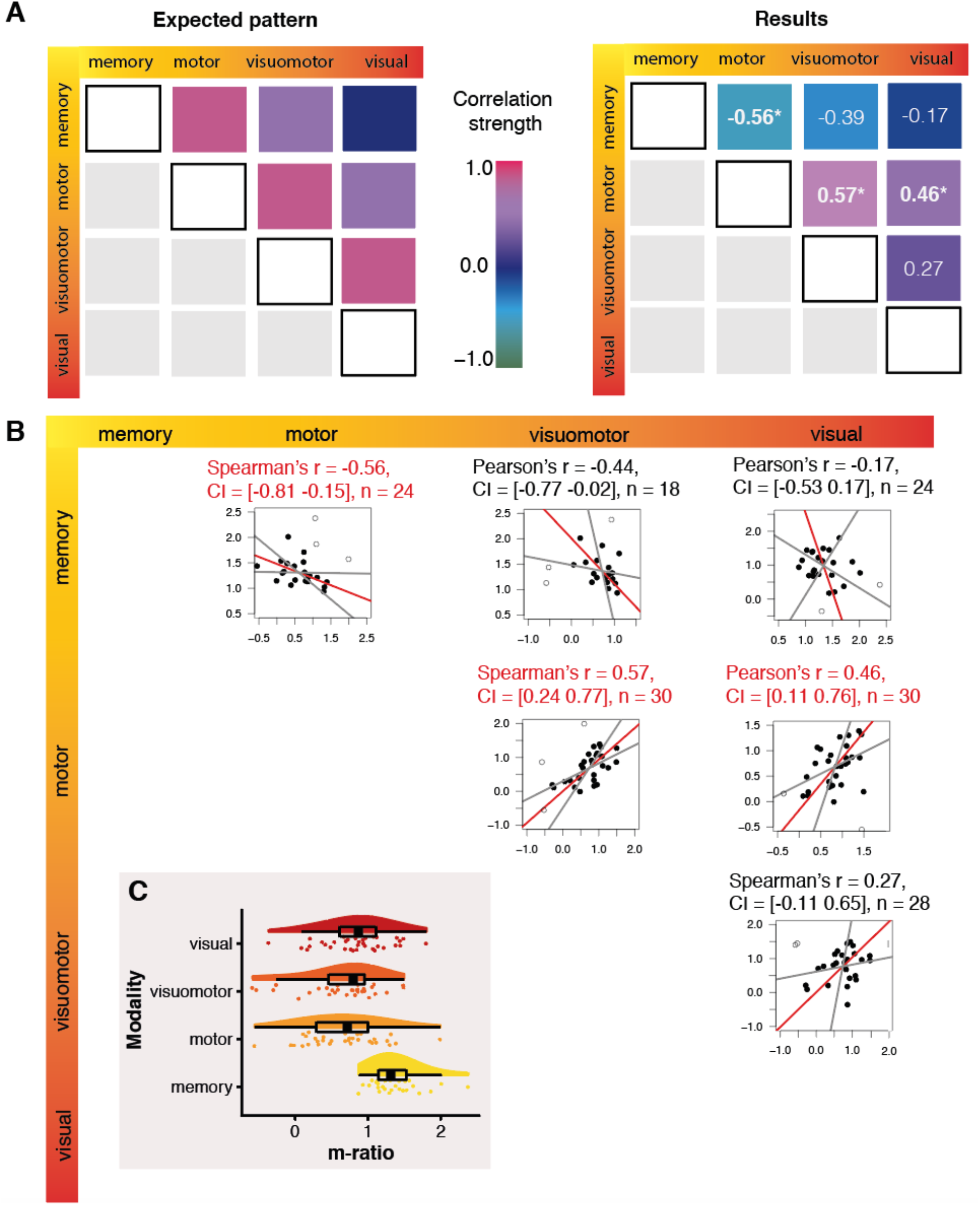
**A. Patterns of correlations expected under the hypothesis of the validity of the internal-external axis of domains, and observed correlations. Colors are schematic. B. Correlations plots.** Those task pairs for which *m-ratios* showed a statistically significant correlation are highlighted with red titles. The grey lines represent the 95% CIs of the parametric slope estimates for the line of best fit, drawn through the centroid of the bivariate distribution. The empty dots in panel E represent outliers that were excluded from the correlation analysis and line of best fit estimation by the robust correlation procedure (Pernet et al., 2013). **C. Metacognitive efficiency (*m-ratios*) for each task**. In these raincloud plots (Allen et al., 2021), split violin plots reflect the probability density of the data, boxplots show the interquartile range (IQR), the thick line represents the median, the whiskers correspond to the ±1.5 IQR, and the dots reflect individual participants’ values.

Importantly, our results are in line with what has been reported in the literature on visual and memory monitoring, serving as an external validation for our methods and general accuracy of our approach. We contribute to the behavioural findings that found no association between memory and visual metacognitive ability (Baird et al., 2013, 2014, 2015; Fitzgerald et al., 2017; Morales et al., 2018). Further, we largely replicated our own previous results where we found relationships between metacognitive ability in visual, motor and visuomotor tasks (Arbuzova et al., 2021), as well as similar correlations between *m-ratios* in a visual and a visuomotor task (Charles et al. 2020). Taken together, our results do not support the idea of grouping metacognitive mechanisms into two broad groups based solely on the distinction between internal and external information. We suggest that this simple dimension is not sufficient to map the relationships between metacognitive domains. Specifically, what we classify as internal information in the memory task and in the motor task might not be so homogenous, and the two tasks might differ in other aspects that have a stronger influence on *m-ratios* than the internal-external dimension.

More generally, our findings also go against the idea of a common underlying factor of metacognition across all modalities (Mazancieux et al., 2020). A similarly heterogeneous picture is observed in other specialized somatic sensations — metacognitive mechanisms of pain (Beck et al., 2019) and interoception (Legrand et al. 2021) were suggested to be different from perceptual metacognitive mechanisms, based, as here, on finding no correlations between estimates of metacognitive efficiency in either task.

Apart from a basic understanding of the mechanisms underlying metacognitive monitoring, there are two main practical motivations to study the domain-generality of metacognitive processes. First, it will allow us to determine the generalizability of research in metacognition: How far can we generalize from findings on one particular domain to all others? So far, the vast majority of metacognitive studies operationalized metacognitive monitoring using perceptual (predominantly visual) tasks (Rahnev et al., 2020). But often the conclusions are extrapolated to metacognition in general (Fleming et al., 2010). The results we present here underline the importance of using a wide and diverse set of tasks to understand relationships between cognitive domains, and not extrapolating from a potentially non-representative sample. A second motivation to probe the domain-generality of metacognitive monitoring is that it may help optimize metacognitive training. The ability to improve the accuracy of metacognitive monitoring through training is appealing, as better metacognitive ability has been related to better learning outcomes. For example, some measures of tendency to engage in metacognitive monitoring have been associated with better academic performance in general (Ohtani & Hisasaka, 2018). Further, better metamemory is related to better use of adaptive strategies (like cognitive offloading, Gilbert et al., 2020; Hu et al., 2019). In order to understand whether metacognitive training should be tailored to the specific domain of interest, or whether training in one domain can transfer to others, it is crucial to understand the relationships between domains. The evidence for cross-domain effects of metacognitive training is scarce and mixed, as some studies have presented evidence that improvements in metacognitive monitoring (but not first-order performance) might transfer between retinotopic locations (Schwiedrzik et al., 2011) and between metacognition of memory and visual perception (Carpenter et al., 2019), the latter has been contested (Rouy et al., 2022).

### Limitations

Our conclusions rest on the assumption that the motor and visuomotor tasks fall in between the two extremes of the internal-external spectrum. It is plausible that this assumption is not valid. More concretely, the motor and visuomotor tasks might not be correctly placed in the middle of the internal-external spectrum. The motor domain contains at least two distinct components: motor commands and proprioceptive signals. Because proprioceptive signals travel through afferent fibers from the body to the brain, we classified them as externally-generated. However, it is often noted that proprioceptive information is processed predominantly unconsciously (Proske & Gandevia, 2012), thus, proprioceptive metacognition might differ from other kinds of perceptual, exteroceptive metacognition. Metacognition of proprioception has not been studied separately. More work is necessary to examine possible relationships between metacognitive mechanisms for these other somatic sensations and motor metacognition, to understand whether motor metacognition can be considered as a strongly perceptual domain, or whether it is a distinct domain, with distinct properties.

Another reason that might explain the pattern of correlations we found is that we may not have controlled for some task differences that play a more important role than previously thought. For example, although similar in appearance, the visual and memory tasks have very different temporal structures. This, in turn, could have different effects on memory and attentional load. In line with this speculation, we found the highest correlation between *m-ratios* in the motor and visuomotor tasks, which are those that differed the least in terms of the task structure, and therefore, presented similar task demands in terms of memory and attentional demands.

To conclude, our results suggest that the internal vs external distinction of modalities to understand the generality of metacognitive ability is not useful as a guiding principle. Other groupings, such as exteroception, interoception, motor and cognitive might provide a more nuanced view of metacognition across different modalities. We also highlight the need to consider the specific features of the tasks when probing the domain-generality of metacognition using correlational analyses.

## Acknowledgements

We would like to thank Anne Felsenheimer for help with data collection, Jorge Morales for making the code for the stimuli publicly available and for advice with the implementation, and Pedro Espinosa for help with data preprocessing and preliminary analysis.

