## Supplementary Materials for "Metacognitive domains are not aligned along a dimension of internal-external information source"

Here, we report the general structure of our main measures: type I performance (*d′*) and mean confidence ratings for all modalities.

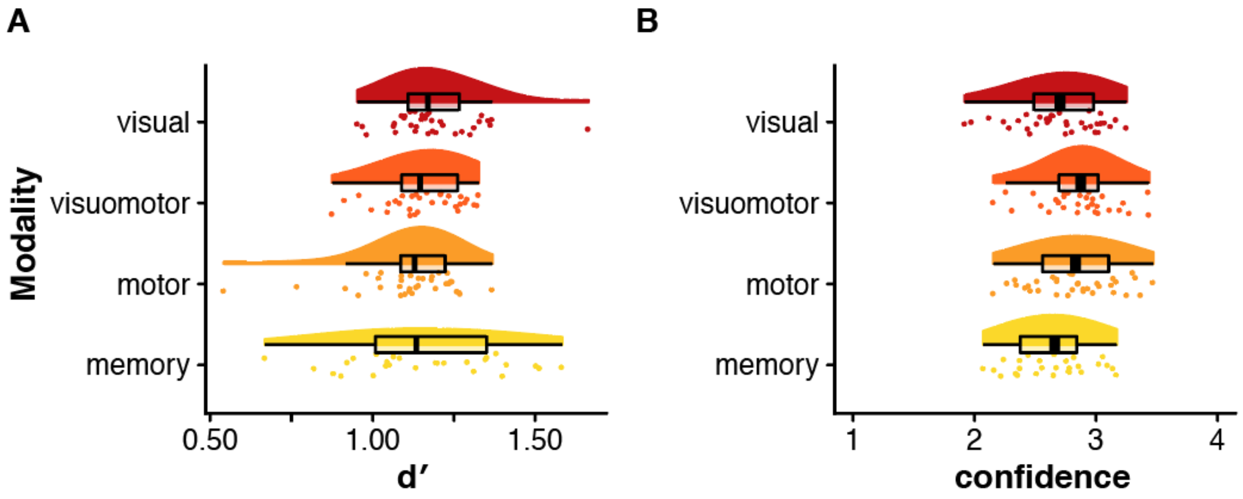


**Figure S1.** Population values for (A) *d′* and (B) mean confidence. In these raincloud plots (Allen et al., 2021), split violin plots reflect the probability density of the data, boxplots show the interquartile range (IQR), the thick line represents the median, the whiskers show ±1.5 IQR, and the dots reflect individual participants' values.

### Type I performance

Since the accuracy of the type I tasks was controlled during the task, either by an online staircase (in visual, visuomotor and motor tasks) or after every block (memory task), we did not expect a difference in *d′*. Indeed, a one-way ANOVA confirms that there was no significant difference in *d′* across tasks: (Greenhouse-Geisser-corrected because of sphericity violations, $F(1.58, 28.50) = 0.90$, $p = .40$, mean (M) confidence and standard deviation (SD): visuomotor – 1.16 (0.12), motor – 1.11 (0.15), visual – 1.19 (0.14), memory – 1.16 (0.23)).

It should be noted that the SD in the memory task is considerably larger than in other tasks (see

Figure S1.A), which can be explained by a coarser online difficulty adjustment and has been observed in the same task previously (Morales et al., 2018).

**Confidence**

We also checked mean confidence ratings across all tasks (Figure S1.B). A one-way ANOVA showed a difference in mean confidence levels (Greenhouse-Geisser-corrected, $F(2.08, 37.48) = 8.72$, $p < .001$, M (SD): visuomotor – 2.86 (0.31), motor – 2.83 (0.34), visual – 2.69 (0.37), memory – 2.64 (0.31)). Bonferroni-corrected, post-hoc pairwise t-tests showed that mean confidence ratings in the memory task were lower than in the motor task ($p = 0.002$), as well as in the visuomotor task ($p < 0.001$). No other pairwise comparison yielded a statistically significant difference.
